# Genome-wide investigation of *WRKY* gene family in *Lagenaria siceraria*

**DOI:** 10.64898/2025.12.30.696992

**Authors:** Yaling Wang, Qing Lai, Min Wang, Hui Zhu, Dafu Ru, Xiaonong Guo

## Abstract

**Background:** A significant family of transcription factors known as *WRKY* genes include many physiological functions and environmental adaptations. However, insufficient information was previously available about the *WRKY* genes in *Lagenaria siceraria*, a crucial crop with substantial economic significance. The recent publication of the whole-genome sequence of *L. siceraria* has allowed us to perform a genome-wide investigation of the organization of the *WRKY* genes in *L. siceraria*.

**Results:** In the present study, 57 *L. siceraria WRKY* (*LsiWRKY*) genes were identified and given new names based on their relative chromosomal distribution. The 57 *LsiWRKYs* were further divided into three major groups and several subgroups based on their structural and phylogenetic properties. Segmentation duplication events have played a major role in the expansion of the *WRKY* gene family in *L. siceraria*. Phylogenetic comparisons of the Group III *WRKY* genes provide valuable insights into the evolutionary characteristics of *WRKY* genes in *L. siceraria*. Additionally, RNA-seq analysis revealed distinct expression pattern of *WRKY* genes across different tissues.

**Conclusions:** This study presents a preliminary analysis of the *WRKY* gene family in *L. siceraria*, including their structural characteristics, evolutionary traits, and tissue-specific expression patterns. The systematic insights provided here serve as a foundation for further functional studies aimed at enhancing *L. siceraria* crops. This knowledge holds promise for improving the cultivation and yield of *L. siceraria*, thereby contributing to agricultural advancements.

## Background

Transcription factors (TFs) are the class of proteins that can interact with other regulatory factors or bind to specific DNA sequences in the promoter regions of genes, thereby regulating the functioning of various genes and thus involved in downstream target genes regulation process (Franco-Zorrilla et al., 2014; Li et al., 2015). As an influential form of TFs, the *WRKY* gene family has been studied for more than two decades since it was first cloned from sweet potato (Ishiguro et al., 1994) and primarily found in single-celled algae and plants. They are named as WRKY because the protein sequence contains several, highly conserved WRKY domains, which include about 60 amino acids. Each and every WRKY protein that has been identified has one or two WRKY domains at the N-terminus, followed by zinc finger motifs at the C-terminus (Li et al., 2015). The classification of WRKY proteins into three broad classes is based on the quantity of WRKY domains and the type of zinc finger sequences (I-III). Members of group I have two WRKY domains and a zinc-finger motif of the C2H2 type, whereas group II and group III only have one WRKY domain and follow it with zinc finger motifs of the C2H2 and C2HC types, respectively (Eulgem et al., 2000). The WRKYs of group II can be further classified into five different subgroups based on their phylogenetic relationship (IIa-e). Through the identification of the W-box core sequence (TTGACC/T) within the promoter region of target genes, *WRKY* transcription factors exhibit exclusive binding to these target genes (Yu et al., 2001).

According to incomplete statistics, more than 14,500 WRKY proteins have been identified from 165 plant species (Jin et al., 2017). The great majority of WRKY protein research have shown that these proteins are engaged in a variety of biological and abiotic stress responses as well as playing important roles in the plant immune system. By way of illustration, increasing the expression of *AtWRKY4* can make plants more susceptible to the biotrophic bacterium *Psudomonas siringae* (Lai et al., 2008). In Cucumis, *LsiWRKY50* plays a positive role in *Pseudoperonospora cubensis* resistance involving multiple signaling pathways (Luan et al., 2019). In comparison to wild-type plants, *OsWRKY47* overexpression can boost rice yield and drought resistance (Raineri et al., 2015). In response to salinity stress, *GmWRKY92*, *GmWRKY144*, and *GmWRKY165* would be positively regulated in soybeans (Song et al., 2016). It has been shown that *OsWRKY11* overexpression can increase tolerance to stress caused by high temperatures (Wu et al., 2009). *VvWRKY30* was proved to confer tolerance to salt stress in *Vitis vinifera* (Zhu et al., 2019). Moreover, WRKY proteins also participate in additional crucial plant processes, such as pollen development (Lei et al., 2017), seed size (Luo et al., 2005; Gu et al., 2017), seed dormancy and germination (Jiang et al., 2009; Xie et al., 2007; Zou et al., 2008), plant development (Johnson et al., 2002; Ishida et al., 2007) and leaf senescence (Ay et al., 2009; Brusslan et al., 2012; Yang et al., 2016). Nearly all WRKY families in angiosperms have undergone significant expansion during evolution due to the substantial involvement of the WRKY family in a variety of physiological activities. For instance, there are at least 70 WRKY proteins in *Arabidopsis* (Eulgem et al., 2000; Dong et al., 2003), 174 in *Glycine max* (Yang et al., 2017b; Yang et al., 2017a) and 109 in rice (Wu et al., 2005).

Although a large number of studies have been published on the *WRKY* gene family, relatively few have investigated the bottle gourd. One of the major crops in the Cucurbitaceae, the bottle gourd (*Lagenaria siceraria*), is a diploid species (2n=2x=22), and it possesses a genome size of 313.4 Mb (Wu et al., 2017). It is believed to have originated in southern Africa and is now widely grown in the tropical and subtropical regions (Wu et al., 2017), particularly in the East Asian countries (Kistler et al., 2014). Due to its beneficial nutritional properties (Loukou et al., 2011) and health properties (Shah et al., 2010), *L. siceraria* has a significant potential for usage in medicines. For instance, it hydrates the skin and decreases edema and knots. It also can be used for food, containers, decorative artefacts or musical instruments (Mashilo et al., 2017). In order to enhance the cold tolerance and disease resistance of other cucurbit crops, bottle gourd has recently emerged as a vital rootstock material for grafting (Davis et al., 2008; King et al., 2008). For instance, the bottle gourd is the recommended rootstock for watermelon, one of the most widely grown fruits in the world because it controls soil-borne diseases and has no impact on fruit quality of the fruit (Davis et al., 2008; Fidan et al., 2016). Therefore, the study of important functional genes in the bottle gourd has aroused substantial interest from researchers. A thorough analysis of the *WRKY* gene family in *L. siceraria* would be crucial due to the significance of the *WRKY* genes in many physiological systems. The recent completion of sequencing of the *L. siceraria* genome provides an opportunity to reveal the organization and evolutionary traits of the *L. siceraria WRKY* gene family at the genome-wide level. In the current work, 57 *L. siceraria WRKY* genes were discovered, and they were divided into three major groups. Comprehensive analyses including the exon-intron organization, motif composition, gene duplication, chromosome distribution, phylogenetic and synteny analysis were also investigated. Our study provides valuable clues to understand the functional characterization of members of the *WRKY* gene family in *L. siceraria*.

## Results

### Identification of the *WRKY* genes in *L. siceraria*

In this study, we systematically investigated the *WRKY* gene family in *L. siceraria*, one of the largest families of plant transcription factors. Initially, 61 putative *WRKY* genes were identified through BLASTP using *Arabidopsis thaliana* genes as references (Table S1). Subsequently, redundant and non-WRKY domain-containing sequences were removed, resulting in the exclusion of specific sequences (Lsi01G000510.1, Lsi05G005250.1, Lsi09G001980.1, Lsi09G010130.1). Ultimately, 57 *WRKY* genes were identified and annotated by validating the presence of WRKY domains using the SMART program. The number of identified *WRKY* genes in *L. siceraria* (Table S2) was comparable to that of other plants, such as *Cucumis Melo* L. (57 members) (Chen et al., 2021), *Cucumis sativus* (57 members) (Ling et al., 2011) and *Capsicum annuum* L. (61 members) (Cheng et al. 2016). These 57 *WRKY* genes were successfully mapped to chromosomes 1–11, and based on their chromosomal locations, they were systematically renamed as *LsiWRKY1* to *LsiWRKY57* (Fig. 1; Table S2).

**Fig. 1.**
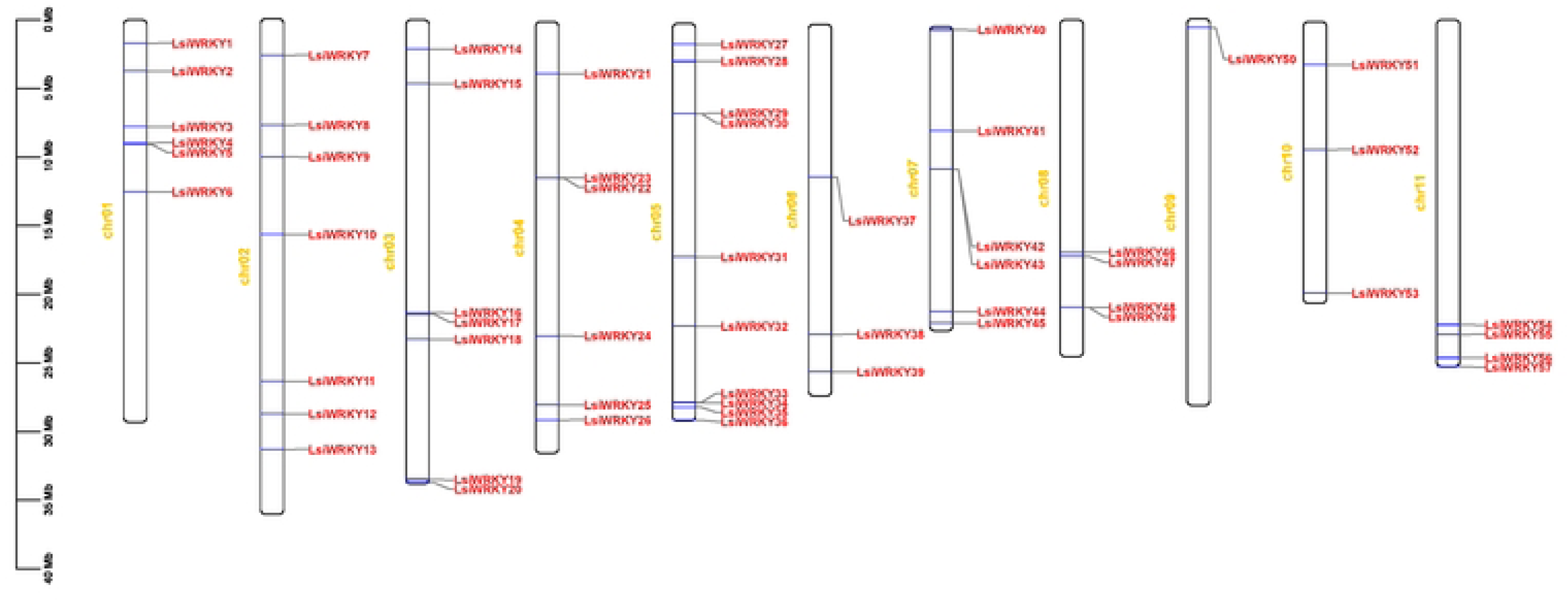
Mapping of the WRKY gene family on *L. siceraria* chromosomes. The size of a chromosome is indicated by its relative length.

Furthermore, we investigated additional essential features of the WRKY proteins, including their protein sequence length, coding sequence (CDS) length, molecular weight (MW), and isoelectric point (pI) (Table S3). Among the 57 LsiWRKY proteins, LsiWRKY04 with 119 amino acids represented the smallest, whereas LsiWRKY44 with 751 amino acids was the largest protein. The molecular weights of these proteins ranged from 13.125 kDa (LsiWRKY04) to 81.409 kDa (LsiWRKY44), indicating significant variability in protein sizes. Additionally, the pI of WRKY proteins spanned from 4.64 (LsiWRKY16) to 9.73 (LsiWRKY38), underscoring the diverse biochemical properties within this gene family. This comprehensive analysis provides a detailed overview of the structural characteristics of the identified LsiWRKY proteins, setting the stage for a deeper understanding of their functional roles in *L. siceraria*.

### Multiple sequence alignment and phylogenetic analysis

In our investigation, we conducted a multiple protein sequence alignment of all 57 LsiWRKY proteins using Muscle software to explore their evolutionary relationships (Fig. S1). Subsequently, using MEGAX software with the neighbour-joining method, a phylogenetic tree was constructed based on the highly conserved WRKY domains of 57 LsiWRKYs and 71 AtWRKYs (Fig. 2). The phylogenetic analysis revealed that the 57 LsiWRKYs could be categorized into three major groups analogous to the grouping in *Arabidopsis* as defined by Eulgem et al. (2000) (Fig. 2). Specifically, 8 LsiWRKY proteins were classified into group I, 39 into group II, and 7 into group III, while 3 remained unclassified (Fig. 3). In group I, the 8 members shared C2H2-type zinc-finger motifs (C-X4-C-X22–23-H-X-H) and possessed both N-terminal and C-terminal WRKY domains. Group II, comprising the majority of LsiWRKYs, was further divided into five subgroups (IIa-IIe). These subgroups exhibited variations in the C2H2-type zinc-finger motifs and contained different numbers of WRKY domains (4 WRKY proteins belonged to IIa, 5 to IIb, 17 to IIc, 7 to IId, and 6 to IIe). Notably, subgroup IIc was the most abundant, mirroring the grouping pattern observed in AtWRKYs. Group III, distinctive due to the zinc-finger motif C2HC: C-X-C-X23-H-X-C, comprised 7 LsiWRKY members, each possessing one WRKY domain.

**Fig. 2.**
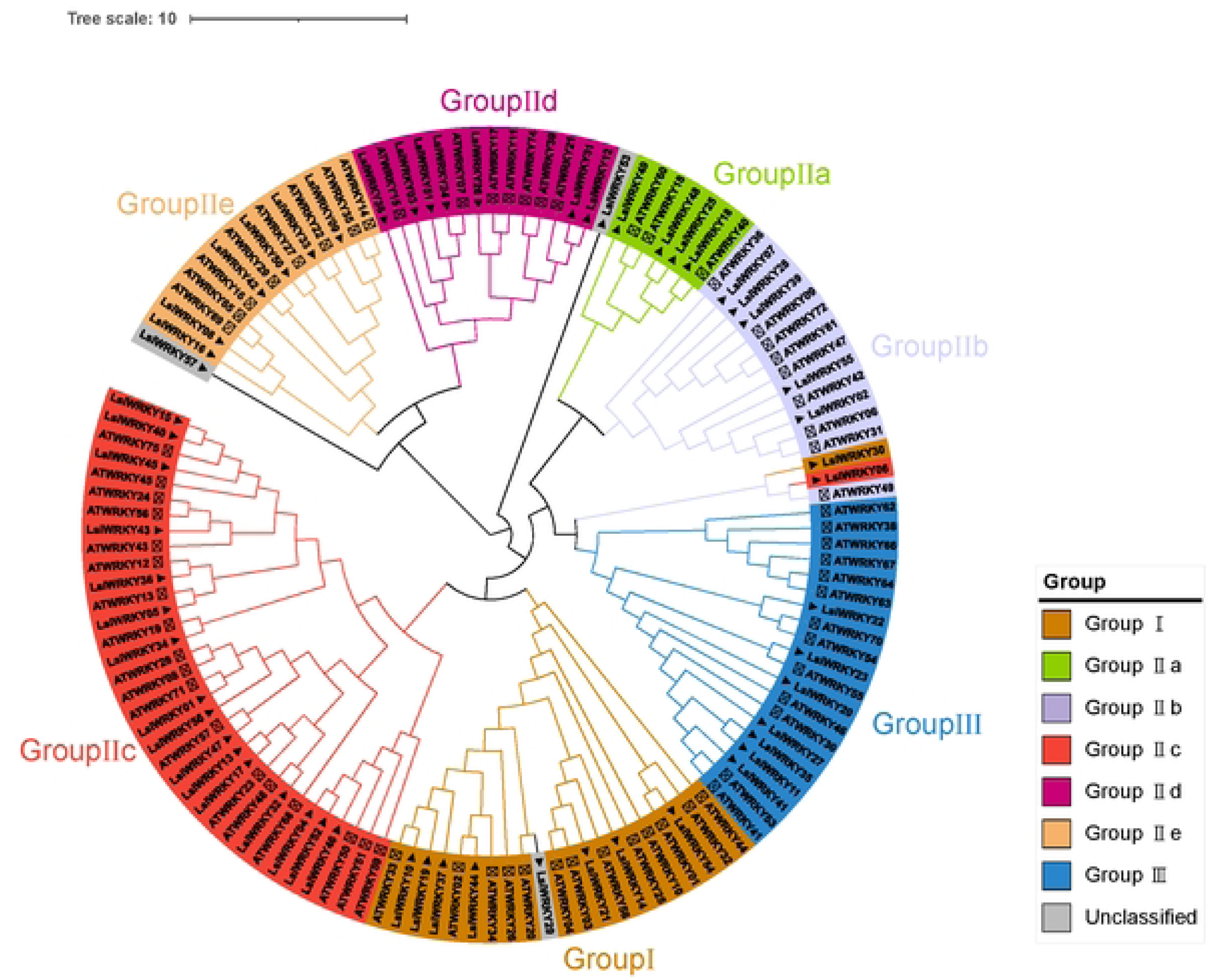
The neighbor joining phylogenetic tree of WRKY family genes of *Arabidopsis thaliana* and *L. siceraria*. Each WRKY group is labeled with different colors.Solid triangles represent *L. siceraria* and hollow triangles represent *Arabidopsis thaliana*.

**Fig. 3.**
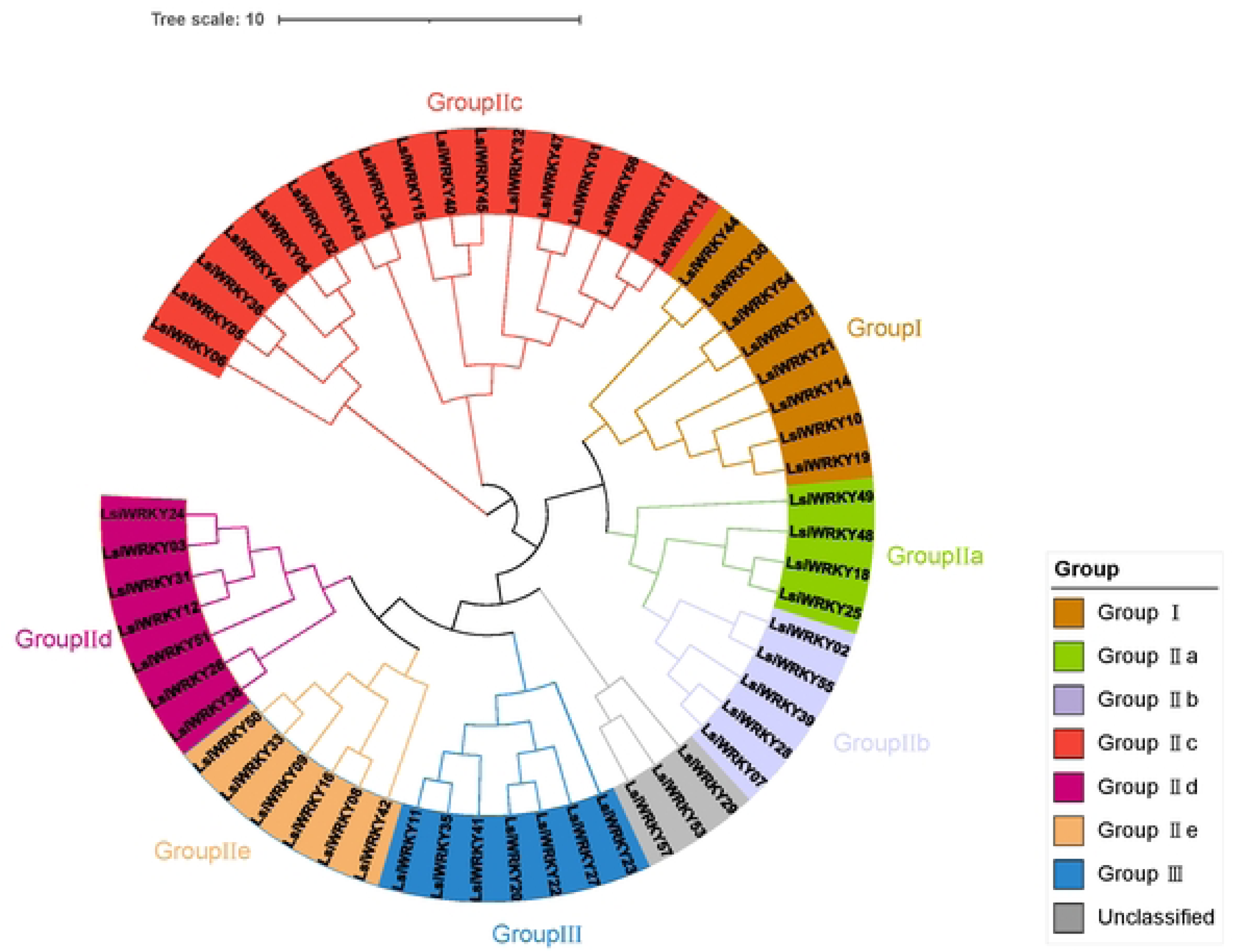
Phylogenetic tree of LsiWRKY proteins in *L. siceraria* using the neighbor joining method by MEGA X.

Interestingly, our analysis identified a potential R-protein WRKY, LsiWRKY34 from group IIc tightly clustered with AtWRKY19, characterized by the presence of the ’leucine-rich repeat’ (LRR) motif, commonly found in resistance (R) proteins (Fig. 3). This suggests a putative role for LsiWRKY34 in *L. siceraria*’s response to biotic or abiotic stresses, akin to the known R-protein WRKYs in *Arabidopsis* (Rinerson et al., 2015; Lobo et al., 2017).

Furthermore, we scrutinized the conserved domain "WRKYGQK", a hallmark of *WRKY* transcription factors, based on the grouping information. While most *LsiWRKYs* exhibited the WRKYGQK variant, indicating Q to K substitutions, three LsiWRKYs (LsiWRKY04, LsiWRKY52, LsiWRKY46) in subgroup IIc displayed WRKYGKK variants (Fig. S1). Additionally, LsiWRKY54 from group I exhibited the mutant WYMRCQM sequence (Fig. S1). These variations, observed mainly in subgroup IIc, mirrored findings in other plant species like peanuts and soybeans, highlighting the sensitivity of WRKY domains in this subgroup to mutations (Song et al., 2016). Furthermore, our analysis revealed certain WRKY domains with substantial sequence variation, leading to their classification into an unclassified group. The origin of these variations could be attributed to potential issues in genomic sequencing or gene prediction programs, warranting further investigation. Overall, these findings shed light on the diversity within the *LsiWRKY* gene family, providing valuable insights into their evolutionary patterns and functional significance.

### Gene structure and motif composition of *L. siceraria* WRKY gene family

The MEME (Multiple EM for Motif Elicitation) tool was employed to reveal the conserved motifs among 57 LsiWRKY proteins in order to more fully characterize the structure of the WRKY domains. In total, ten conserved motifs, labeled as Motifs 1 through 10, were identified (Fig. 4). These motifs varied in width, spanning 21 to 44 amino acids residues, and were represented by distinct colored boxes (Fig. 4). Among these motifs, Motifs 1 and 3 encoded the conserved WRKY domain, while Motifs 2 and 4 encoded the conserved zinc finger structure. All LsiWRKY proteins have one or two WRKY motifs (Fig. 5). Additionally, conserved motifs (Motifs 4–10) were identified in various LsiWRKY proteins. Furthermore, motif 1 was prevalent in almost all LsiWRKYs except for three unclassified genes and *LsiWRKY04*, *LsiWRKY05*, *LsiWRKY30*, *LsiWRKY52* (Fig. 5). Each LsiWRKY protein harbored at least two conserved motifs, and some contained as many as six motifs (Fig. 5).

**Fig. 4.**
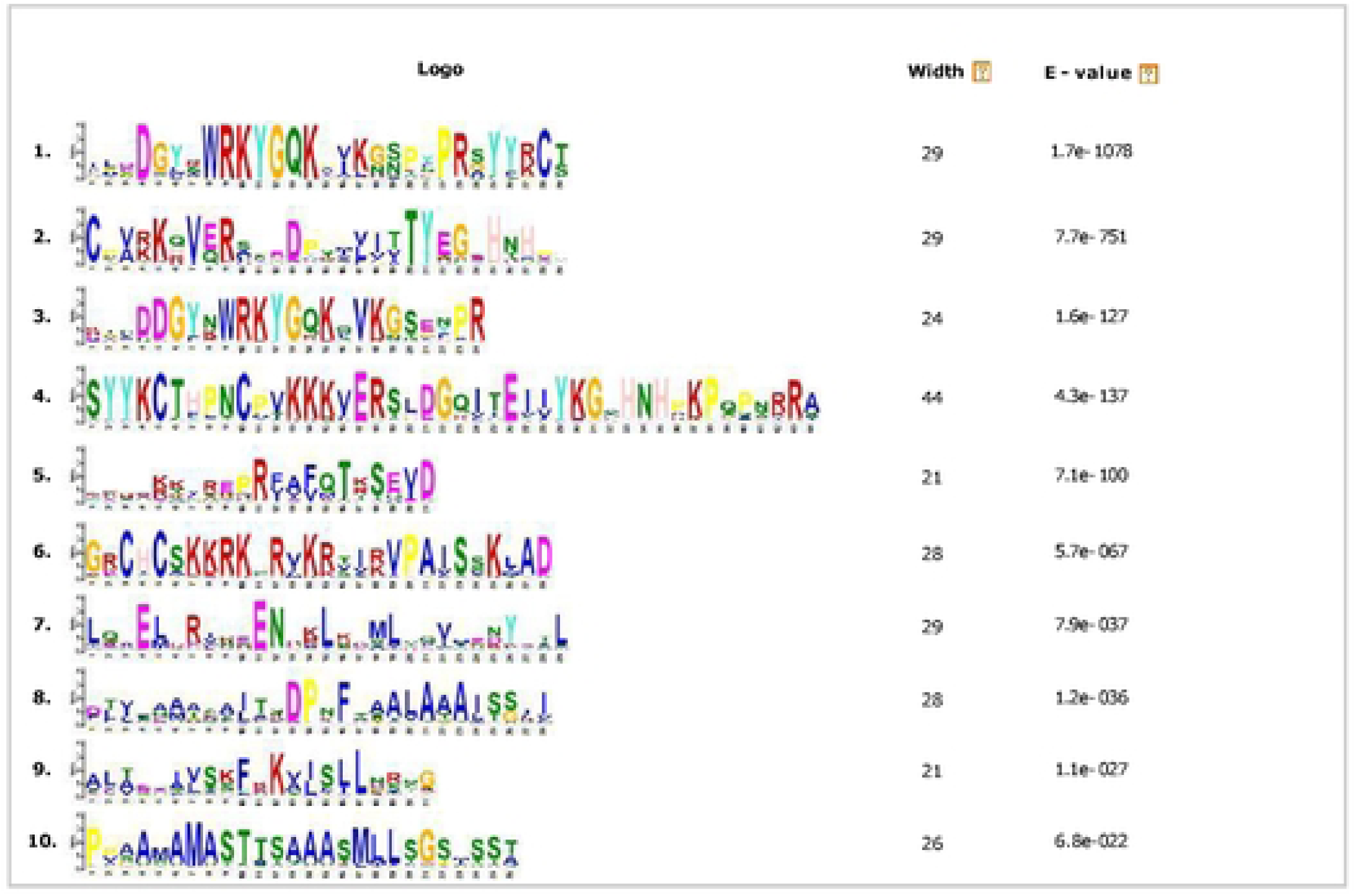
Schematic diagram of conserved motif of WRKY protein in *L. siceraria* including the motif logos, consensus sequence widths in aa, and E-values.

**Fig. 5.**
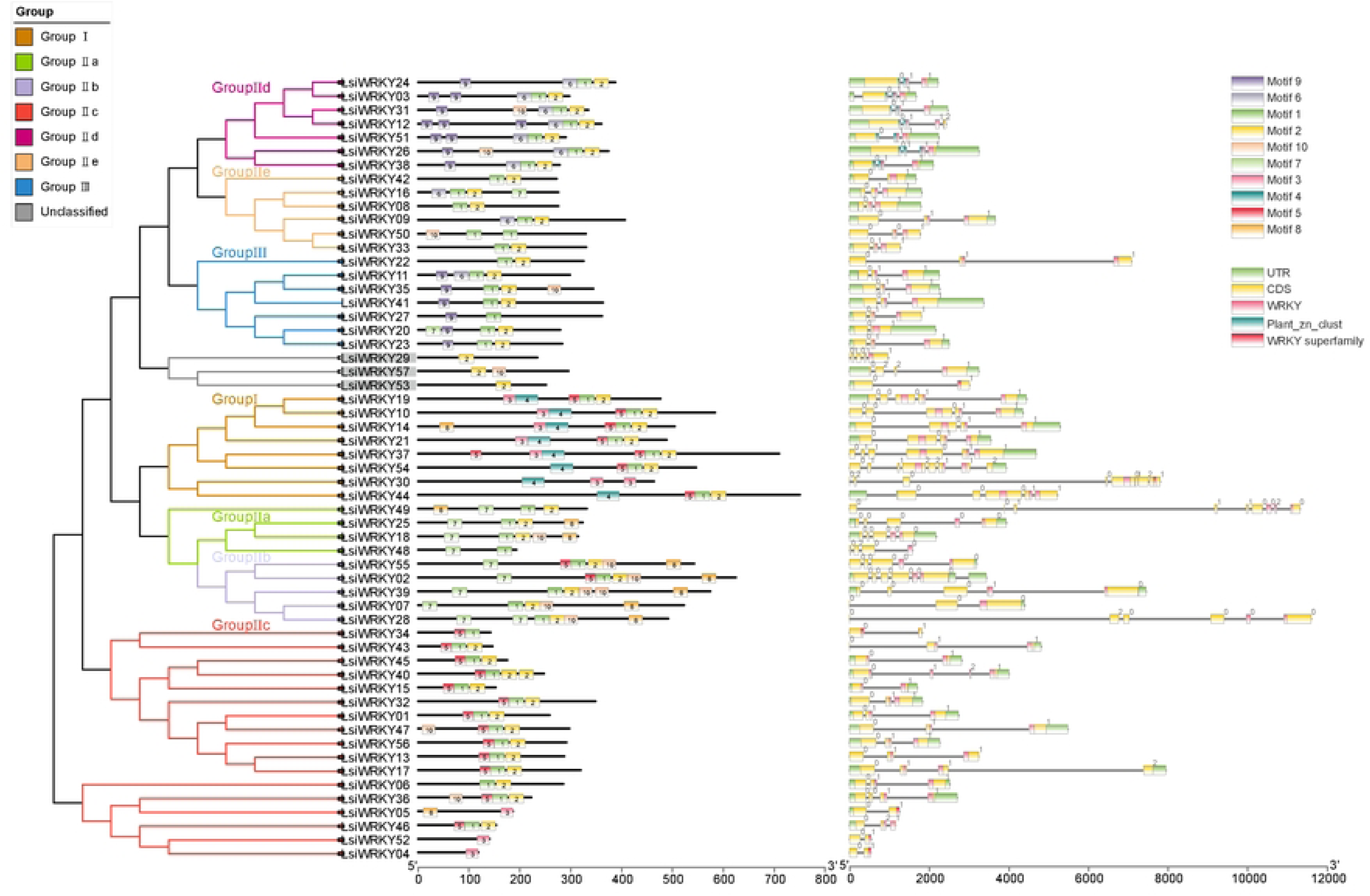
Comprehensive schmatic diagram of phylogenetic clustering,conserved protein motifs, LsiWRKYs gene structure. Left panel: the phylogenetic tree was constructed from the WRKY domain sequences of LsiWRKY proteins. Different colors represent different categories. Middle panel: the motifs are represented by different colored boxes with corresponding numbers. Right panel: gene structure of LsiWRKY. Untranslated 5 ′- and 3 ′-regions, exons, introns, WRKY domains, Plant-zn-clust and WRKY superfamily are indicated by green boxes, yellow boxes, black lines, pink boxes, blue-green boxes and red boxes, respectively. Intron phases 0, 1, and 2 are indicated by numbers 0, 1 and 2, respectively.

Distinct distributions of conserved motifs were observed across different LsiWRKY groupings. For example, group I LsiWRKYs exhibited 3 to 6 motifs (Motifs 1, 2, 3, 4, and 5), with each member in this group possessing at least one of Motifs 1 and 3, as well as at least one of Motifs 2 and 4 (Fig. 5). Notably, Motif 4 was unique to the group I LsiWRKY proteins (Fig. 5). Motif 9 was exclusively present in Group III and subgroup IId, while Group I and subgroups IIb and IIc contained either Motif 4 or Motif 8 (Fig. 5). Subgroups IIa and IIb predominantly featured Motif 7 (Fig. 5). The conservation of motif types within the same group indicated similar functionalities among members.

Studies have shown that the intron-exon structure of multiple gene families plays an important role in plant evolution (Wang et al., 2023; Safder et al., 2021). In order to further understand the structural features of the *WRKY* family in *L. siceraria*, we investigated the exon-intron structures of identified *LsiWRKY* gene. It is obviously that all *LsiWRKY* genes exhibit two to six exons, with none having only one exon (Fig. 5). Generally, genes within the same group share a similar structure, as highlighted in brown for group I members (Fig. 5). Intriguingly, each WRKY domain in *LsiWRKY* genes possessed an intron, except for specific genes such as *LsiWRKY42*, *20*, *29*, *53*, *52*, *57*, *48*, *7*, *5*, and *4*, and *34* (Fig. 5). Intron distributions and phase coincided with the alignment of the *LsiWRKY* genes clusters. V-type intron (phase-0 intron) were found in group IIa and II, while R-type intron (phase-1 intron) akin to those in rice and *Arabidopsis*, were prevalent in other groups (group I, IIc, IId, IIe and III) (Rinerson et al., 2015), with N-terminal WRKY domains of group I lacking introns (Fig. 5).

### Chromosomal distribution of LsiWRKY genes

Based on the information about the location of the genes on the chromosomes, we determined the positional distribution of the 57 *LsiWRKYs* on 11 chromosomes in the *L. siceraria* genome. From the outputs of the MEME motif analysis, a schematic representation of the structure of all LsiWRKY proteins was constructed. With the exception of Motifs 1 and 2, which are broadly dispersed LsiWRKY domains, the distribution of LsiWRKY members within the same group is visualized across all *L. siceraria* chromosomes (Fig. 1; Fig. 5). We found that the distribution of *WRKY* genes on each chromosome was not uniform and dense (Fig. 1). Notably, chromosome 9 harbored only one *LsiWRKY* gene, whereas chromosome 5 exhibited the largest number of *LsiWRKY* genes (n=10), which accounted for 17.5% of all *LsiWRKY* genes. In addition, chromosomes 1, 4 and 7 contain the same number of *LsiWRKY* genes, each hosting 6 LsiWRKY loci. Interestingly, several regions of high *LsiWRKY* gene density were found on some chromosomes, including 1, 3, and 5, suggesting potential *WRKY* gene hotspots in the genome.

### Expression of *WRKY* gene family in different tissues of *L. siceraria*

Previous studies have indicated substantial variation in the expression levels and roles of *WRKY* genes across different tissue (Wang et al., 2019; Fan et al., 2018). To explore the expression patterns of different *WRKY* genes in *L. siceraria* tissues, we analyzed 10 sample replicates (three fruits, two stems, three leaves and two roots) using R (v 4.2.3). Our analysis revealed diverse expression patterns among different WRKY gene groups.

In group I, several genes exhibited high expression across all tissues. Notably, *LsiWRKY10* displayed elevated expression levels in both fruits and stems, while *LsiWRKY19* and *LsiWRKY37* exhibited lower expression in leaves (Fig. 6). Subgroup IIa genes, with the exception of *LsiWRKY48* and *LsiWRKY49*, were highly expressed in all tissues. Conversely, subgroup IIb genes displayed relatively lower expression in all tissues. Subgroup IIc genes exhibited notable functional differentiation, resulting in significant expression variations across tissues. Subgroup IId genes, except for *LsiWRKY31*, showed high expression levels in all tissues, indicating a limited role in leaf development. Subgroups IIe and IIa demonstrated similar expression patterns, being expressed in all tissues. Group III featured high expression of *LsiWRKY35* and *LsiWRKY11*, whereas *LsiWRKY20* exhibited high expression in roots and stems, with other genes displaying lower expression. Genes in the unclassified group exhibited high expression levels in all four tissues, except for *LsiWRKY53* (Fig. 6).

**Fig. 6.**
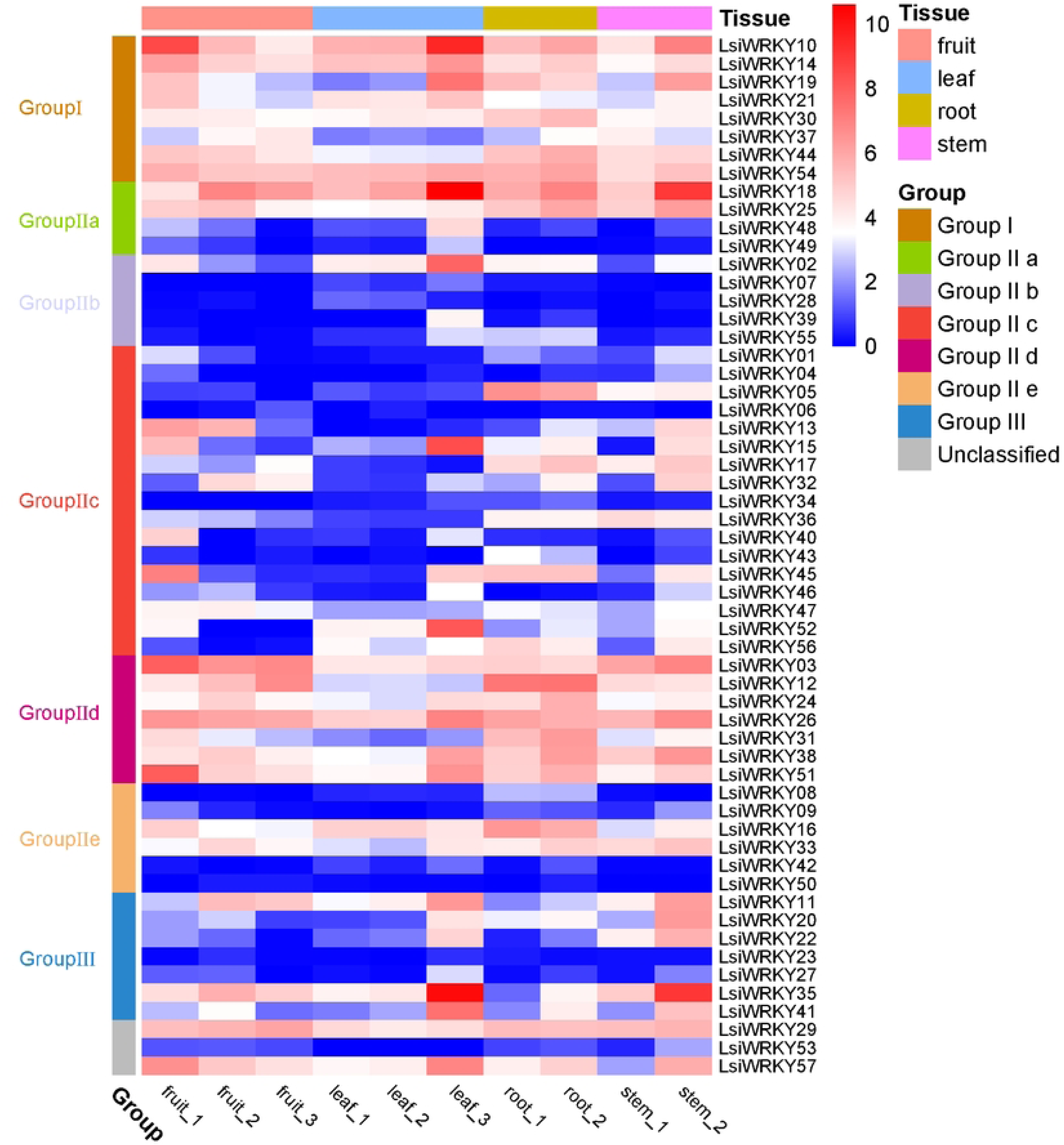
The heatmap of expression levels of 57 WRKY genes in *L. siceraria*.

## Discussion

*WRKY* genes are a family of transcription factors found in all plant species as they play crucial roles in regulating plant development and responses to biotic and abiotic stresses. In this study, we used a genome-wide search which identified 57 putative *WRKY* genes in *L. siceraria*, a plant species of economic importance. Through phylogenetic and structural analyses, we categorized the *WRKY* genes into three groups with several subgroups on the basis of phylogenies and the basic structure of the WRKY domains. Notably, we found that the Group I WRKY proteins in *L. siceraria* retained both WRKY domains and did not undergo any domain loss events during evolution, in contrast to what has been observed in other plant species. These findings shed light on the evolutionary history of the *WRKY* gene family in *L. siceraria*, and provide a basis for further investigations into the functional diversity and regulation of these genes in response to different environmental stimuli.

Previous studies have suggested that N-terminal WRKY domains exhibit weak DNA-binding activity and are more variable during evolution. Among the different subgroups of WRKY, Group I, which contains two WRKY domains, is considered to be the most ancient member that occurred during the evolution of WRKY. The WRKYs in subgroup IIa and IIb are believed to have originated from an algal single WRKY domain or from the other Group I derived lineage (Rinerson et al., 2015; Waqas et al., 2019). Members of subgroup IIc evolved from WRKYs in subgroup II that lacked N-terminal domains. However, the origin of each type of WRKY protein in *L. siceraria* is currently unknown. Although WRKY domains are strongly conserved among WRKY proteins, LsiWRKY proteins exhibit some degree of structural divergence. The heptapeptide WRKYGQK is the typical domain of the WRKY family, but three variants of this domain, including WRKYGKK, have been identified in several LsiWRKY proteins in subgroup IIc. Similar variants of the WRKY domain have also been found in other plant species (Li et al., 2015; Guo et al., 2014; Yue et al., 2016; Zou et al., 2016), suggesting that these variants may confer multiple biological functions to *WRKY* gene family (Wu et al., 2017).

Subgroup IIa contains seven *LsiWRKY* genes and is phylogenetically closer to subgroup IIb than to subgroups IIc. This classification is supported by the fact that the WRKY domains of subgroup IIa and subgroup IIb maintain a similar consensus structure (Fig. S1). In addition, the results of the dual phylogeny (Fig. 2) also showed anomalies compared to the results of the single species phylogeny (Fig. 3), particularly with regard to the positions of *LsiWRKY30*, *LsiWRKY06*, and *ATWRKY49* which clustered together. *LsiWRKY30* should belong to Group Ⅰ, *LsiWRKY06* to subgroup Ⅱc, and *ATWRKY49* to subgroup Ⅱc (Wang et al., 2014; Chen et al., 2020). Based on the close relationship between *LsiWRKY06* and *ATWRKY49* and both of which belong to subgroup Ⅱc, it appears that *LsiWRKY30* was misclassified as subgroup Ⅱc. This may be due to the introduction of *ATWRKY49* in the single-species phylogenetic tree resulting in sequence variation in the WRKY structural domain of LsiWRKY30 and LsiWRKY06 that was not evident during the tree construction, and the phylogenetic software used was not precise enough. Future studies should conduct to further differentiate these sequences, which will help to delineate more precise phylogenetic relationships. Moreover, from an overall perspective, the WRKY domain of ATWRKY49 forms a separate cluster of its own, which indicates a high degree of divergence between this domain and other members of the subgroup Ⅱc. The differentiation between LsiWRKY06 and LsiWRKY30 was also evident, although to a lesser extent. Therefore, the results suggest that subgroup Ⅱc and Group Ⅰ are closely related genetically and eventually diverged into two different subgroups (Fig. 2). It is hypothesized that ATWRKY49 is closely related to subgroup IIc and Group I. Furthermore, the sequence alignment results showed that the WRKY domains of LsiWRKY29, LsiWRKY53, and LsiWRKY57 contained a large degree of variation, which placed them in an unclassified subgroup not previously known in the WRKY family (Fig. 3; Fig. 2). It is worth noting that gene annotation errors may occur (which are rare) due to problems with genome sequencing or gene prediction software, and further validation is needed to confirm the identity and function of these WRKY gene.

In addition, we have identified LsiWRKY34 as a chimeric protein that contains both R-protein and WRKY domains. Chimeric proteins with both domains have been reported in other plant species and have been implicated in plant defense against diseases and stresses. For example, in barley, WRKY1/2 inhibited basal defense, but when the Avra10 effector was present, R protein MLA10 and WRKY1/2 interacted in the cell nucleus to suppress the effect of WRKY1/2 on basal defense and enhance disease resistance (Shen et al., 2007). Similarly, in *Arabidopsis*, the ATWRKY genes encodes an NBS-LRR-WRKY protein that acts as a chimeric protein, with the WRKY domain exhibiting DNA-binding activity (Noutoshi et al., 2005). The presence of R protein-WRKY chimeric proteins in *L. siceraria* suggests a possible role in plant resistance to disease or other stresses, as has been observed in *A. thaliana* and barley. However, further investigation is needed to determine the putative functions of these chimeric proteins in *L. siceraria*. Overall, the results of this study have important implications for understanding the environmental resistance in plants and could potentially lead to the development of new strategies for improving plant productivity and stress resistance.

Our RNA-seq expression analysis of *LsiWRKY* genes revealed their presence in all examined tissues, displaying diverse and distinct expression patterns. Generally, most *LsiWRKYs* displayed relatively high abundance in stems and roots, whereas expression levels were comparatively lower in fruits and leaves. However, specific genes exhibited high expression specifically in fruits and leaves. Furthermore, except for the subgroup IIc, where gene expression patterns varied significantly, genes in other subgroups exhibited similar expression patterns, suggesting potential functional redundancy. Conversely, *LsiWRKYs* with diverse expression patterns likely fulfill different biological functions in plant growth and development. These results lay the groundwork for in-depth analysis of individual *WRKY* gene expressions in *L. siceraria*, shedding light on the intricate regulatory mechanisms within this gene family.

## Conclusions

A comprehensive analysis of the *WRKY* gene family in *L. siceraria* was performed in the present study. Fifty-seven full-length *WRKY* genes were characterized and further classified into three main groups, with strongly similar exon-intron structures and motif compositions within the same groups and subgroups. Through phylogenetic comparison of *WRKY* genes from several different plant species and tissue-specific expression analysis, we gained valuable clues about the evolutionary characteristics of *L. siceraria WRKY* genes. Notably, our identification of a chimeric R-protein-WRKY protein in *L. siceraria* suggests a potential role in disease resistance or stress responses, although further investigation is warranted to fully elucidate its functional significance. The findings provide a solid foundation for future research, offering promising avenues for enhancing agronomic traits and bolstering environmental resistance in this pivotal crop species. Furthermore, our study underscores the broader importance of the *WRKY* gene family in plant biology, shedding light on their specific roles in *L. siceraria*. Overall, this research not only deepens our knowledge of *WRKY* genes but also opens new doors for harnessing their potential in crop improvement, emphasizing their vital role in the context of plant biology.

## Methods

### Gene identification

To identify candidate datasets for LsiWRKYs, the *L. siceraria* whole genome protein database was analyzed using BLASTP (v 2.14.0+; E-value 1e-10; 71 AtWRKYs protein sequences were used as a query), where AtWRKYs protein sequences were downloaded from the TAIR database (https://www.arabidopsis.org). All candidate LsiWRKY protein sequences were eventually characterized for structural domains through the SMART plugin (http://smart.embl-heidelberg.de/) of the TBtools platform (https://github.com/CJ-Chen/TBtools) to test the WRKY conserved domains for integrity, incomplete sequences were excluded from the dataset, and redundant sequences were further manually removed. Finally, the Expasy ProtParam tool (http://us.expasy.org/tools/protparam.html) was used to calculate the biophysical properties of the *LsiWRKY*s protein sequences, including sequence length, molecular weight and protein isoelectric point.

### Gene structure analysis and chromosome Location

A schematic diagram of the exon-intron organization of *L. siceraria WRKY* genes was constructed using the online Gene Structure Display Server (GSDS) (http://gsds.cbi.pku.edu.cn/; accessed time 2023.08.30), in which exon position information was obtained from the genome’s annotated ‘gff’ file. After further information on the distribution of *LsiWRKY* genes on chromosomes was obtained from the genome annotation files, TBtools was utilized to show the distribution of their positions on 11 chromosomes.

### Multiple sequence alignment and MEME analysis

Multiple WRKY protein sequences of *L. siceraria* were first aligned using MUSCLE (v 5.1) with default parameters, and the final results of the alignment were visualized using the Genedoc (v 2.7) software and presented in Fig S1. Next, to identify conserved motifs in *L. siceraria* proteins, the online MEME program (Multiple Expectation Maximization for Motif Elicitation; http://meme-suite.org/tools/meme/2023.08.30) was used to identify conserved motifs in the 57 identified *L. siceraria* WRKY protein sequences. The maximum value of the basic search was set to 10, and the optimum width of each motif was limited to 21-44 amino acid residues.

### Phylogenetic analysis

To elucidate the evolutionary relationships within the *WRKY* gene family of *L. siceraria*, a robust methodology was employed. Firstly, multi-protein sequence alignments were performed between *A. thaliana* and *L. siceraria* and within *L. siceraria* itself. Subsequently, phylogenetic trees were constructed using both neighbor-joining (NJ) and maximum likelihood (ML) algorithms in the MEGA X software.

For the NJ analysis, the Dayhoff substitution matrix (PAM250) was used, and the reliability of the constructed trees was verified through 1000 bootstrap replicates, ensuring robustness and accuracy in the inferred relationships.

In the construction of the ML tree, the best-fit model (JTT+G) was determined from 59 amino acid substitution models using the modelfinder tool in MEGA X. This careful model selection process ensured the appropriateness of the chosen model for the dataset. Following model selection, protein sequence information was integrated, leading to the development of a tree for *L. siceraria’*s WRKY proteins, utilizing the ML algorithm. The resulting ML tree was visually represented using the iTOL tool (https://itol.embl.de/), enhancing the clarity and accessibility of the evolutionary insights. Additionally, the classification of the *LsiWRKY* gene family was meticulously conducted based on the results derived from the phylogenetic tree analysis and the identification of conserved domains. This rigorous approach ensured a comprehensive understanding of the evolutionary dynamics and structural features within the *WRKY* gene family of *L. siceraria*.

### Transcriptome data analyses

Transcriptomic data, comprising three leaf tissues, three fruit seeds, two root tissues, and two stem tissues, were downloaded from National Center for Biotechnology Information (NCBI, https://www.ncbi.nlm.nih.gov/). A total of ten transcriptomic data were included in the analysis.

Prior to analysis, the data underwent rigorous filtering using trim_galore (v 1.18; https://www.bioinformatics.babraham.ac.uk/projects/trim_galore/). Clean reads were then mapped to the *L. siceraria* genome using HISAT2 (v 2.2.1). The resultant data were assembled using featureCounts (v 2.0.6). For quantification, the expression value of each gene was measured in Transcripts Per Kilobase per Million mapped reads (TPM) and calculated using the R (v 4.2.3). R package Heatmap was employed to create an expression level heatmap for different tissues based on log2(TPM+1) data.

## Acknowledgements

This study was supported by Northwest Minzu University Talent Introduction Project (Z2101707), the National Natural Science Foundation of China (grant no. 32001085) and Fundamental Research Funds for Central Universities (grant no. lzujbky-2020-34).

## Supplementary Material

Fig. S1. Schematic diagram of multiple-sequence alignment of conserved WRKY domains. Top panel:the conserved N-terminal LsiWRKY domains of different groups in L. siceraria.And highlight for the variant YMRC sequence. Bottom panel: the conserved C-terminal LsiWRKY domains of different groups in L. siceraria. That belong to the same group are clustered together and marked with different colors. The conserved amino acids are highlight with homochromatic background.

Fig. S2. LsiWRKY proteins domains prediction.

Table S1. List of the 61 putative WRKY genes were initially identified.

Table S2. List of the 57 LsiWRKY genes identified in this study.

Table S3. Physicochemical property of LsiWRKY proteins and grouping.

Table S4. Results of predicting the conserved domain of LsiWRKY genes using the CD-search tool.

Table S5. ten transcriptome data and their download sources.

